# A learned score function improves the power of mass spectrometry database search

**DOI:** 10.1101/2024.01.26.577425

**Authors:** Varun Ananth, Justin Sanders, Melih Yilmaz, Sewoong Oh, William Stafford Noble

**Author notes:** Equal contributions (Varun Ananth, Justin Sanders).

## Abstract

One of the core problems in the analysis of protein tandem mass spectrometry data is the peptide assignment problem: determining, for each observed spectrum, the peptide sequence that was responsible for generating the spectrum. Two primary classes of methods are used to solve this problem: database search and *de novo* peptide sequencing. State-of-the-art methods for *de novo* sequencing employ machine learning methods, whereas most database search engines use hand-designed score functions to evaluate the quality of a match between an observed spectrum and a candidate peptide from the database. We hypothesize that machine learning models for *de novo* sequencing implicitly learn a score function that captures the relationship between peptides and spectra, and thus may be re-purposed as a score function for database search. Because this score function is trained from massive amounts of mass spectrometry data, it could potentially outperform existing, hand-designed database search tools. To test this hypothesis, we re-engineered Casanovo, which has been shown to provide state-of-the-art *de novo* sequencing capabilities, to assign scores to given peptide-spectrum pairs. We then evaluated the statistical power of this Casanovo score function, Casanovo-DB, to detect peptides on a benchmark of three mass spectrometry runs from three different species. Our results show that, at a 1% peptide-level false discovery rate threshold, Casanovo-DB outperforms existing hand-designed score functions by 35% to 88%. In addition, we show that re-scoring with the Percolator post-processor benefits Casanovo-DB more than other score functions, further increasing the number of detected peptides.

## 1 Introduction

Tandem mass spectrometry is a high-throughput protein content analysis method that can be used to investigate a broad range of biological phenomena, including cellular mechanisms, disease progression, protein alterations, and protein-protein interactions. Each tandem mass spectrometry experiment produces as output a collection of spectra, where each spectrum ideally was generated by a distinct peptide sequence. Hence, a central analysis challenge is the peptide assignment problem, wherein each observed spectrum is assigned a corresponding peptide sequence. Correctly matching peptides to their respective spectra allows for accurate identification and quantification of peptides in a sample, which is critical for the aforementioned downstream applications of tandem mass spectrometry.

The standard approach for solving the peptide detection problem is database search. Database search algorithms solve the peptide detection problem by searching an observed MS2 spectrum against a pre- defined database of peptides. The query spectrum is compared, using some scoring function, to a theoretical spectrum for each database peptide within a given *m/z* tolerance, and the top-scoring peptide-spectrum match (PSM) is returned. Currently, all widely used database search engines use hand-designed score functions to evaluate the similarity between an observed spectrum and each candidate peptide from the database. The first such score function was the XCorr score, implemented in SEQUEST [1] and used by popular search engines such as Comet [2] and Tide [3], but many others have been designed subsequently (reviewed in [4]). While differing somewhat in implementation details, all such score functions take the same general approach of measuring the number of shared fragment ions between the observed mass spectrum and an idealized theoretical spectrum for a given peptide. However, these score functions have been shown to be under powered [5], poorly calibrated [6], and unable to fully capture the complex fragmentation patterns governing the relationship between peptides and their fragment ion spectra [7, 8]. These limitations may lead to systematic under-detection of peptides in proteomics datasets, leaving valuable insights hidden in the data.

Traditional database search is feasible only when working with data with a corresponding, well-characterized proteome, such as human or model organism data. In some settings, including metaproteomics, antibody sequencing, paleoproteomics, and immunopeptidomics, such a database is not available. In these settings, the alternative solution to the peptide assignment problem is *de novo* peptide sequencing, in which the peptide sequence is inferred directly from the observed spectrum without the use of a peptide database. Recently, deep learning methods have achieved state-of-the-art performance on this *de novo* sequencing task [9–12]. These models are trained in a supervised fashion to predict a peptide sequence given an observed spectrum.

Here, we hypothesize that when trained on the *de novo* sequencing task, these deep models implicitly learn the relationship between spectra and their generating peptides. Thus, we propose using the confidence score assigned to predictions from a *de novo* sequencing model as a score function for database search. Because such models are trained from labeled peptide-spectrum pairs first identified via database search, this proposed score function bootstraps from existing hand-derived score functions, but further benefits from the massive quantities of annotated mass spectra available in public data repositories. Here, we show that leveraging this wealth of available data to learn a better score function can increase the power and sensitivity of existing database search algorithms. Specifically, a powerful score function is one that detects many peptides at a given false discovery rate (FDR), as estimated by a standard statistical model known as “target-decoy competition” [13]. Here, we use a pre-trained, state-of-the-art *de novo* sequencing model, Casanovo [12], to assign scores to given peptide-spectrum pairs during database search. Our results suggest that, at a 1% FDR threshold, the Casanovo-DB score function consistently yields a larger number of detected peptides than existing score functions, including XCorr [1], the X!Tandem Hyperscore [14], and Andromeda [15]. In particular, we identify between 31–40% more peptides than XCorr, 52–69% more peptides than Hyperscore, and 88–102% more peptides than Andromeda score on datasets from *E. coli*, yeast and human samples.

## 2 Methods

### 2.1 Casanovo-DB score function

Our proposed learned score function is derived from the deep learning model Casanovo [12]. Casanovo is trained on *∼*30 million high confidence PSMs to solve the *de novo* sequencing problem, in which the amino acid sequence of a peptide is inferred directly from an observed mass spectrum. Casanovo treats the *de novo* sequencing problem as a sequence-to-sequence translation task, where the spectrum is represented as a sequence of peaks and the output is the predicted sequence of amino acids. The model consists of a transformer architecture popular in the field of natural-language processing [16], with a transformer encoder learning an in-context representation of the input mass spectrum and a transformer decoder to predict the next amino acid in the peptide sequence given the spectrum representation and the previously predicted amino acids. Inference with Casanovo is auto-regressive: the model predicts a peptide sequence one amino acid at a time, at each step choosing the most likely next amino acid given the current predicted prefix.

Here, we propose to instead use Casanovo to score only the peptides present in the protein database during database search. To score a given peptide spectrum match using Casanovo, rather than letting the model predict each amino acid, we instead input the true sequence of the peptide into the model and record the score assigned to each amino acid in the sequence. Thus, for a given input spectrum and a peptide sequence *s*_1_, *s*_2_, …, *s*_*n*_, we obtain as output from Casanovo a list of predictions *x*_1_, *x*_2_, …, *x*_*n*_, where each *x*_*i*_ represents the models estimated likelihood that *s*_*i*_ is the next amino acid in the peptide given the spectrum and the prefix *s*_1_, …, *s*_*i−*1_. To combine these per-amino-acid scores into a single PSM score, we take the geometric mean:

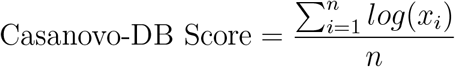

We choose to use the geometric mean of amino acid scores, rather than the arithmetic mean used by Casanovo, in order to more severely punish peptides where Casanovo is uncertain for some amino acids in the sequence. Unlike the *de novo* sequencing setting, in which only the highest scoring peptide for each spectrum is important, here we want the model to also accurately assign low scores to peptides which are not a good match for the given spectra. Thus, we found that penalizing low amino acid confidence scores more severely by using the geometric mean gave better experimental results.

Conceptually, this scoring procedure is analogous to the “teacher forcing” that is employed during Casanovo training, and allows us to extract a measure of what the trained Casanovo model estimates is the plausibility that the given spectrum was generated by the candidate peptide. Higher scores indicate a greater confidence that the match is valid.

To run a full database search using the Casanovo-DB score function, the model scores each observed spectrum with respect to all candidate peptides in the database that fall within a user-specified mass range, specified in units of parts-per-million (ppm). This is a standard procedure for performing mass spectrometry database search. Casanovo produces as output an mzTab file [17] in which, for each spectrum, the top-scoring candidate peptide is reported along with its score.

### 2.2 Existing score functions

We selected three widely used search engines with distinct score functions to compare our method to: Tide search [3] which uses the XCorr score function, SAGE [18] which uses the Hyperscore score function, and MaxQuant [15] which uses the Andromeda score function.

XCorr is the oldest score function, proposed in the very first database search algorithm, SEQUEST [1], and still used by the Tide and Comet search engines [2]. XCorr scores a peptide-spectrum match as follows. First, the mass spectrum *u* is binned at 1.0005079 Da/charge resolution and normalized. Second, a theoretical spectrum *v* is constructed for the peptide, containing constant intensity peaks for the b-, y-, and a- fragment ions along with neutral losses of ammonia and water. Finally, the similarity of *u* and *v* is calculated as

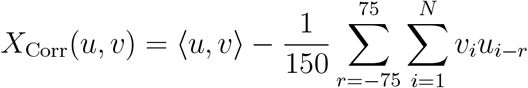

Thus, XCorr is the cross-correlation between the observed and theoretical spectra, giving a measure of their dot product similarity normalized by the background similarity observed between the two vectors at various bin offsets.

The Hyperscore takes a more probabilistic approach to scoring PSMs, comparing the number of observed fragment ions for the given peptide compared to the number expected purely by chance under a hyper- geometric distribution. Specifically, the Hyperscore counts the number of matched b-ions *n*_*b*_ and y-ions *n*_*y*_, along with their unit-normalized intensities *I*_*b*_ and *I*_*y*_, and computes:

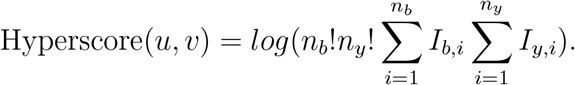

The Andromeda score takes a similar approach, but uses the binomial distribution to assign significance to the number of observed fragment ions. The given mass spectrum is pre-processed by keeping only the *q* most intense peaks per 100 Da/charge window, and a theoretical spectrum *u* is constructed for the peptide containing the same set of fragment ions as used by XCorr. The Andromeda score is then calculated as

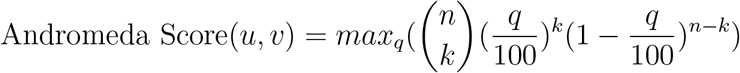

where *n* is the number of peaks in the theoretical spectra for the peptide and *k* is the number of matching peaks observed in the spectrum.

### 2.3 Performance measures

We evaluate each search engine by measuring the number of distinct peptides that are detected from a given dataset at a false discovery rate (FDR) threshold of 1%. FDR is estimated using a standard procedure, known as “target-decoy competition” (TDC) [13]. Here, we use a “double competition” variant of TDC to estimate FDR at the peptide level [19] The procedure consists of the following steps:

1. Digest the proteins in the given database to peptides using a pre-specified set of enzymatic digestion rules. The resulting peptides are referred to as “targets.”
2. Shuffle each unique target peptide to create a corresponding decoy peptide.
3. Score each observed spectrum with respect to all of the target and decoy peptides whose masses lie within a specified range of the spectrum’s associated precursor mass.
4. For each spectrum, retain the single, top-scoring target or decoy peptide. We refer to each spectrum and its associated peptide as a peptide-spectrum match (PSM). This is the first competition, because the target and decoy peptides compete to be assigned to the spectrum.
5. For each peptide that is matched to at least one spectrum, retain only the top-scoring PSM for that peptide.
6. For each target peptide that is matched to at least one spectrum, compete it against the corresponding decoy peptide, retaining whichever of the two PSMs has a better score. Any target or decoy that is not matched to any spectrum automatically loses the competition.
7. Rank the remaining peptides by score, from largest (i.e., best) to smallest. At each position *k* in the ranked list, estimate the FDR as min 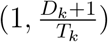 where *D*_*k*_ (respectively, *T*_*k*_) is the number of decoy peptides (respectively, targets) with rank smaller than *k*.
8. Select the largest value of *k* such that 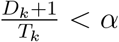, for some specified random matching rate *α*. In this work, we used *α* = 0.01.

We use Crema [20] to perform decoy-based FDR estimation on the PSM rankings provided by running a database search on the aforementioned .mgf files and corresponding .fasta files.

As an auxiliary performance measure, we also compute for each score function and dataset the “target match percentage” (TMP) [21], which is defined as the proportion of spectra that are assigned to target peptide. Formally, the TMP is 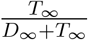. The TMP is complementary to the number of peptides accepted at 1% FDR, because the TMP does not require that scores be calibrated across spectra. By independently evaluating the ranking for each spectrum, a PSM score function can perform well according to the TMP measure even if the top-scoring scores assigned to two different spectra are scaled differently and cannot be directly compared to one another. For two TMP values to be comparable, they need to be computed on the same set of spectra. Due to differences in implementation details, different search engines filter out different spectra based on certain quality control metrics, and thus do not report the top scoring PSM for all input spectra. This makes it impossible to directly compare the TMP for the results. Therefore, we perform a TMP comparison only between Casanovo-DB and XCorr, which assign PSMs to all spectra in the dataset making TMP values comparable.

### 2.4 Benchmarking data

To evaluate the performance of the four score functions, we downloaded a publicly available dataset (ProteomeXchange ID: PXD028735) containing mass spectrometry runs for the organisms *Escherichia coli* (*E. coli*), *Homo sapiens* (human), and *Saccharomyces cerevisiae* (yeast) which is commonly used as a benchmark for mass spectrometry analysis pipelines [22]. Because these are MS/MS runs that were generated post-2019, we guarantee that there is no train/test leakage for Casanovo-DB, since Casanovo was trained on mass spectra generated and published prior to 2019 [12]. All of the selected data was generated using an Orbitrap instrument, run in data-dependent acquisition mode. The resulting .raw files were converted to .mgf files using MSConvert [23]. Peak picking for MS levels 1–2 using the “Vendor” algorithm was employed to reduce noise. The human, yeast, and *E. coli* .fasta files used in all database searches were downloaded from UniProt on 11/6/23, 4:30 PM.

#### 2.4.1 Database search parameters

To ensure a fair comparison for our performance benchmark, we ran each search engine with equivalent parameters wherever possible. All searches employed a single, static modification, carbamidomethylation of cysteine (C+57.02146). Variable modifications included methionine oxidation (M+15.994915), asparagine and glutamine deamidation (N+0.984016, Q+0.984016), as well as four N-terminal modifications (X+42.010565, X+43.005814, X-17.026549, X+25.980265). The cleavage rule was trypsin with proline suppression ([RK]— {P}), and we set the allowed maximum missed cleavages to 1. Finally, the precursor ion mass tolerance and fragment ion mass tolerance were *±*20 ppm each. In some cases, it was not possible to full standardize all parameters. Other than the above settings, we chose to leave all other options as their default values in all other search engines. For example, the maximum number of variable modifications per peptide was kept to search engine defaults.

#### 2.4.2 Percolator re-scoring

We used Percolator version 3.06.01, as implemented in Crux version 4.1-5c7d0d1-2023-11-14 [24], to perform semi-supervised PSM re-scoring. The feature list provided to Percolator for all search engines was minimal, consisting only of score, precursor m/z, and charge. All other options were left as defaults. This minimal use of Percolator simply calibrates all score functions with respect to charge and *m/z*. After re-ranking PSMs, Percolator automatically performs peptide level FDR control, and generates a peptide-level report assigning a q-value to each peptide detection.

## 3 Results

### 3.1 Casanovo-DB is more powerful than existing score functions

In our first experiment, we aimed to compare the statistical power of four score functions to detect peptides from mass spectrometry data while controlling the false discovery rate. We carried out this analysis on data from three different species, *E. coli*, yeast, and human, and we used target-decoy competition to control the FDR. In each case, we find that the learned Casanovo-DB score function detects more peptides across a range of FDR thresholds in all three species (Figure 1). Specifically, at the standard 1% FDR threshold, we observe an average 88% improvement over the Andromeda score (*E. coli*: 88%, yeast: 75%, human: 102%), 57% improvement over the Hyperscore (*E. coli*: 69%, yeast: 50%, human: 52%), and 35% improvement over XCorr (*E. coli*: 31%, yeast: 35%, human: 40%).

**Figure 1:**
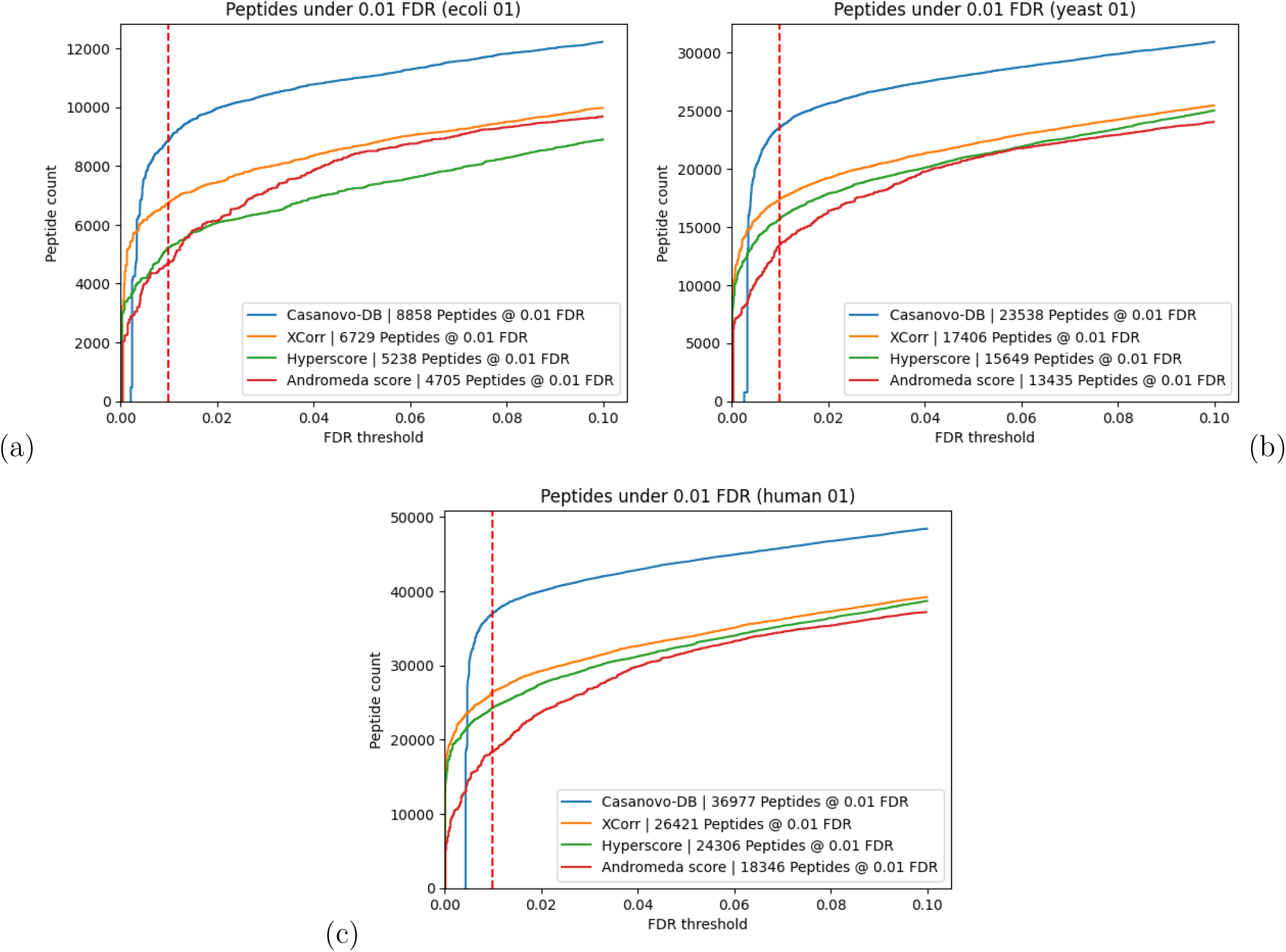
Each figure plots the number of peptides detected as a function of FDR threshold for the (a) *E. coli*, (b) yeast and (c) human datasets. In each plot, the series correspond to different score functions, and the 1% FDR threshold is highlighted with a red dashed line.

We also use a complementary measure, the target match percentage (TMP), to evaluate the quality of each score function. The TMP is the proportion of spectra whose top-scoring peptide is a target. This measure does not take into account how well calibrated a given score is across spectra. Hence, achieving a high TMP score is necessary but not sufficient to achieving good statistical power. We observe that Casanovo-DB again outperforms the next most powerful score function, matching more spectra to target peptides than XCorr in all three species (Table 1). This result implies that at least some of Casanovo-DB’s strong performance in Figure 1 is due to its ability to rank the generating peptide above other candidate peptides, irrespective of score calibration across spectra. Note that it is not possible to calculate the TMP for Andromeda and Hyperscore because the corresponding search engines include filters that prevent scores from being assigned to some spectra.

**Table 1:** Target match percentage (TMP) comparison between Casanovo-DB and Tide for the *E. coli*, human, and yeast datasets. Casanovo-DB outperforms Tide for all organisms.

| Organism | Casanovo-DB | XCorr |
| --- | --- | --- |
| *E. coli* | **0.707** | 0.679 |
| human | **0.819** | 0.776 |
| yeast | **0.760** | 0.725 |

An important component of Casanovo-DB is the use of the geometric, rather than arithmetic, mean of Casanovo’s per-amino-acid score. In the context of *de novo* sequencing, we have observed that these two methods for combining scores perform very similarly. However, in the database search setting, switching from arithmetic to geometric mean dramatically improves the performance of Casanovo-DB. For example, at a 1% FDR threshold, the number of peptides detected increases by 21% for the *E. coli* data, 6% for the yeast data, and 14% for the human data. We hypothesize that this difference arises due to the increased need for calibration in the database search setting. In the *de novo* setting, Casanovo can search for a relatively high quality peptide for every given spectrum, whereas in the database search setting, Casanovo is sometimes forced to score a spectrum with respect to a small set of candidates, none of which truly generated the spectrum. Hence, the database search setting requires comparing high quality PSMs with low quality PSMs, making calibration more important.

### 3.2 Percolator re-scoring benefits Casanovo-DB more than other score functions

In practice, database search tools are rarely used on their own. Instead, their output is typically post-processed using a tool such as Percolator [25], which employs a semi-supervised machine learning algorithm to re-score PSMs based on additional features of the peptide and spectrum. This re-scoring serves to address the calibration issues exhibited by many score functions by adjusting PSM scores based on confounding factors such as peptide length, precursor m/z, and charge. Because Casanovo-DB scores are derived from a deep learning model that was not specifically optimized to produce well calibrated scores, we hypothesized that Percolator re-scoring would benefit Casanovo-DB search results to a greater extent than other score functions.

To test this hypothesis, we repeated our comparison of score functions, but including a Percolator post-processing step in each case. The results (Figure 2) support our hypothesis, with Percolator further increasing the number of additional peptides detected by Casanovo-DB at 0.01 FDR. Percolator improves the number of identifications for Casanovo-DB by an average of 9%, compared to improvements of 5% for XCorr, 8% for Hyperscore and 21% for Andromeda. Andromeda’s strong improvement from Percolator suggests that it may also be poorly calibrated across spectra. Overall, these results further support the idea that Casanovo-DB’s power comes primarily from its ability to rank the generating peptide at the top of the list of candidates. Therefore, when coupled with Percolator, which improves the calibration of those scores across spectra, we achieve even better statistical power.

**Figure 2:**
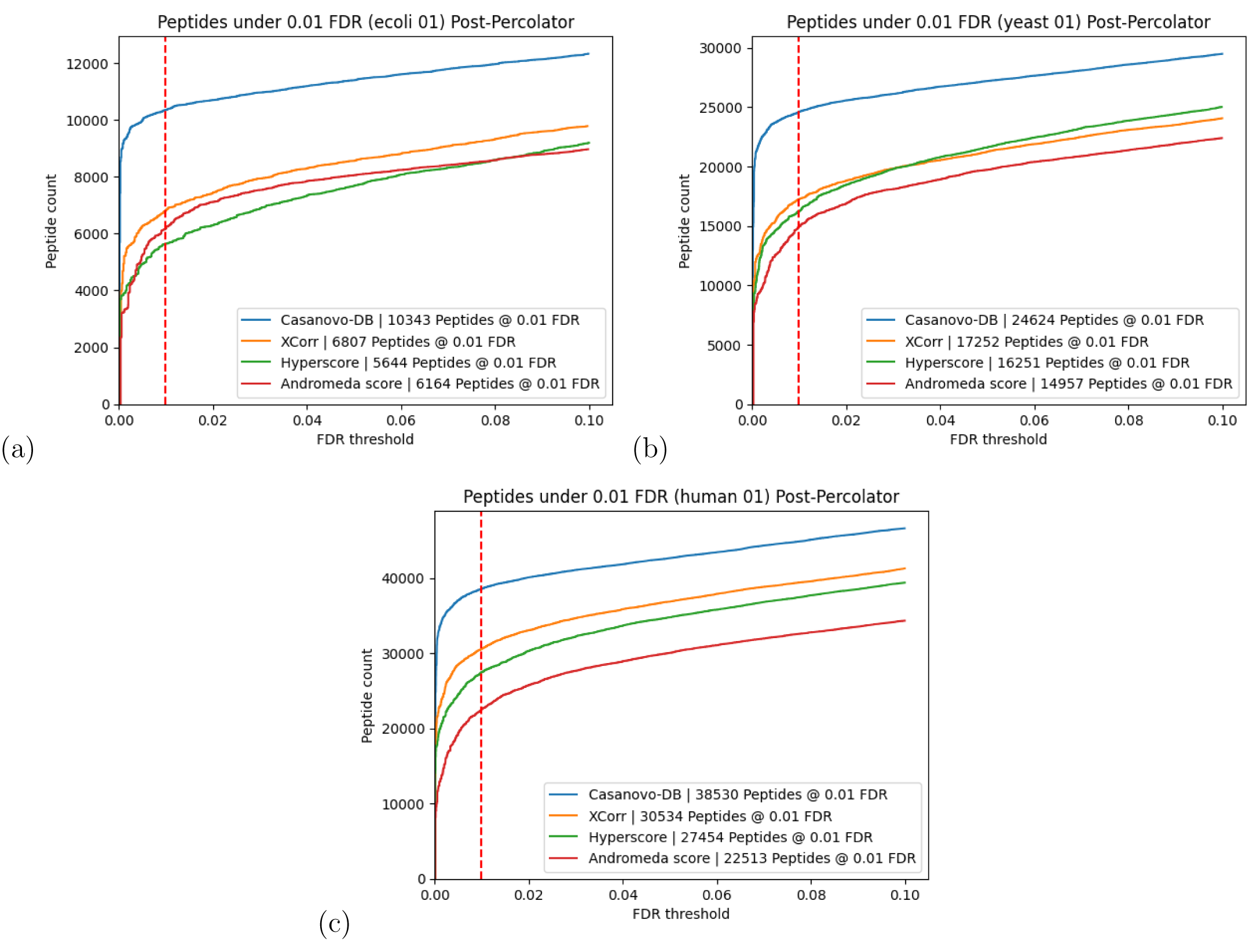
Each figure plots the number of peptides detected after Percolator post-processing as a function of FDR threshold for the (a) *E. coli*, (b) yeast and (c) human datasets. In each plot, the series correspond to different score functions, and the 1% FDR threshold is highlighted with a red dashed line.

### 3.3 Casanovo-DB peptide detections are consistent with existing score functions

Next we compared the set of peptide detections from Casanovo-DB to those obtained by the other three score functions. The existing score functions we benchmark against are widely used by the mass spectrometry community and give trusted results, so any new score function should produce results consistent with existing methods. Reassuringly, we observe a very high overlap between Casanovo-DB and the other score functions, with very few peptides reproducibly detected by other score functions but not by Casanovo-DB (Figure 3). On the other hand, Casanovo-DB detects a large number of peptides (9442) that are not detected by other score functions at 1% FDR. Further investigation revealed that 6812 (72.1%) of the peptides detected only by Casanovo-DB are successfully detected by at least one other search engine when we relax the FDR threshold to 10%. Overall, these results indicate that using Casanovo-DB as a score function for database search has the potential to yield new discoveries while also reproducing the findings obtained from existing methods.

**Figure 3:**
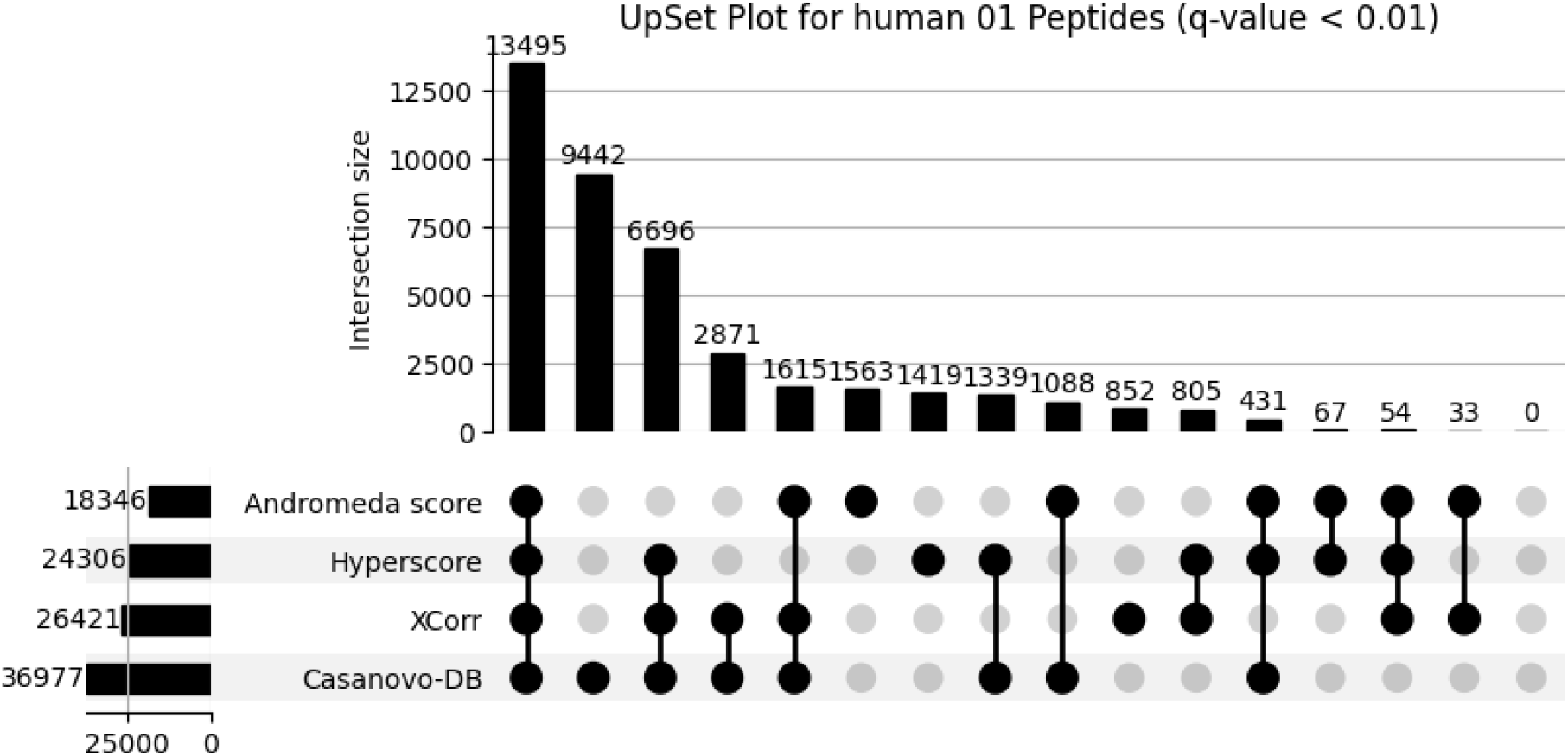
An upset plot showing the overlap in peptide detections at 1% FDR between Casanovo-DB, Tide, SAGE, and MaxQuant on the human dataset.

Finally, to further understand what new peptides are being detected by Casanovo-DB, we investigated how the performance of each search engine varies by peptide mass. Specifically, we segregated spectra according to their associated precursor *m/z* values to find those in the top quartile (704–1389 *m/z*) and bottom quartile (350–467 *m/z*). We then performed TDC separately on each subset. We find that all of the search engines tend to make mistakes on shorter peptides at a much higher rate, while performing significantly better on longer peptides (Figure 4). This analysis explains why, before Percolator re-scoring, Casanovo-DB exhibits worse performance than existing score functions at very small FDR thresholds: there are a handful of very short or highly charged decoys which Casanovo-DB assigns a high PSM score to. This also explains why Percolator re-scoring improves our model by such a large margin, because Percolator is able to re-calibrate Casanovo-DB’s overconfident predictions on low *m/z* precursors.

**Figure 4:**
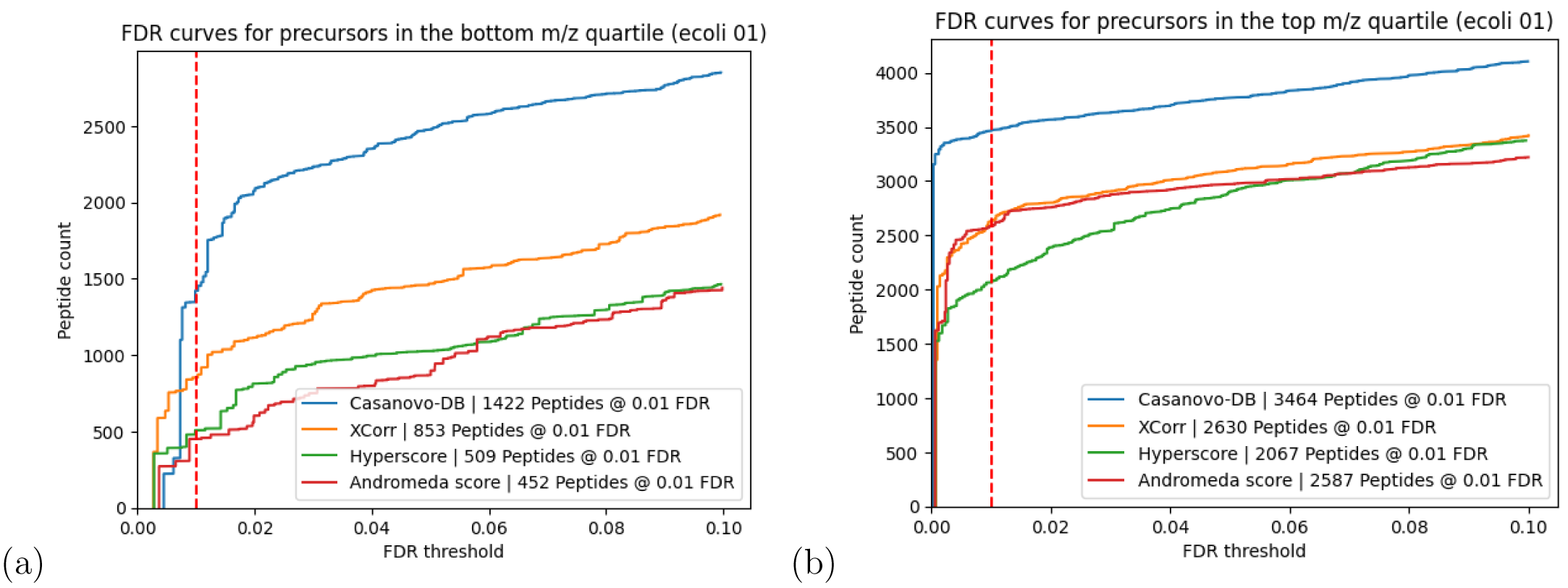
Each figure plots the number of peptides detected as a function of FDR threshold for the *E. coli* dataset, broken down by whether the precursor had an *m/z* in (a) the bottom quartile range of 350–467 *m/z* or (b) the top quartile range of 704–1389 *m/z*. In each plot, the series correspond to different score functions, and the 1% FDR threshold is highlighted with a red dashed line. Casanovo-DB performs much worse on low *m/z* precursors compared to those with high *m/z*, exemplifying the calibration problems which are resolved by Percolator.

## 4 Discussion

Our results show that a trained *de novo* sequencing model can be directly repurposed to carry out mass spectrometry database search, yielding a greater number of detected peptides while controlling for false discoveries. Tandem mass spectrometry assays are commonly used to understand disease progression, accelerate drug development, and deepen our understanding of various proteomes. Database search is the workhorse method for identifying peptide and protein concentrations from tandem mass spectrometry experiments carried out in hundreds of labs around the world. Therefore, increasing the statistical power of scoring functions in this workflow could lead to important discoveries in downstream analyses.

The performance of Casanovo-DB is particularly noteworthy because it is, to our knowledge, the first score function to be learned purely from massive amounts of labeled mass spectrometry data. Hand designed score functions like XCorr struggle to capture the complexities of how peptides fragment in a mass spectrometer, and while post-processing tools can address calibration, they do not improve the power of the score function itself. The success of Casanovo-DB is also especially exciting given the fact that the original task that Casanovo was trained on (*de novo sequencing)* is only tangentially related to the problem of mass spectrometry database search. This suggests that fine-tuning a Casanovo model directly on the task of maximizing peptide detections during database search may further improve detection power.

While there are still a number of steps necessary to make Casanovo-DB usable in practice, in particular addressing the computational cost and hardware requirements inherent to deploying large deep learning models, our results are a compelling first step towards using a fully learned score function to improve the power of database search. With further optimization and fine-tuning, we see the potential to improve standard mass spectrometry analysis pipelines with a learned PSM score function that outperforms traditional hand-designed score functions by a large margin.

